# Advanced eMAGE for highly efficient combinatorial editing of a stable genome

**DOI:** 10.1101/2020.08.30.256743

**Authors:** Zhuobin Liang, Eli Metzner, Farren J. Isaacs

## Abstract

Eukaryotic multiplex genome engineering (eMAGE) offers a powerful tool to generate precise combinatorial genome modifications in *Saccharomyces cerevisiae*. We optimize the design of synthetic oligonucleotides and enrichment of edited populations to increase editing frequencies up to 90%, reduce workflow time by 40%, and engineer a tunable mismatch repair system to lower the rate of spontaneous mutations ∼17-fold. These advances transiently evade genome maintenance to introduce multiple edits at high efficiencies in a stable genetic background, expanding utility of eMAGE in eukaryotic cells.

## Main

Genome editing has empowered researchers to elucidate causal links between genotype and phenotype, enact targeted genetic modifications to reprogram cellular behavior, and design organisms with synthetic genomes^1^. Conventional genome editing technologies use programmable endonucleases, including zinc-finger nucleases (ZFNs), transcription activator-like effector nucleases (TALENs) and CRISPR-Cas nucleases, to generate site-specific DNA double-strand breaks (DSBs) in the genome. Subsequent DSB repair by endogenous machinery can introduce mutagenic gene disruptions via non-homologous end joining (NHEJ) or precise modifications via homology directed repair (HDR)^2^. Recent advances in expression of multiple guide RNAs and orthologous Cas nucleases have enabled multiplex CRISPR-Cas editing of 3-10 sites simultaneously^3^. In addition, several methods derived from CRISPR explore fusion of nuclease-deficient Cas proteins to a variety of DNA effectors, such as deaminases^4, 5^, error-prone polymerases^6^, transposases^7, 8^ and reverse transcriptases^9^, to develop DSB-independent, alternative genome editing tools (*e.g*., Base editing^4, 5^ and Prime editing^9^). However, for applications requiring generation of multisite (>10) combinatorial genomic alterations with base-pair (bp)-level precision, conventional or modified CRISPR-based genome editing systems are inadequate due to inherent limitations^10^, such as off-target effects, cytotoxicity, editing accuracy, and multiplexing ceiling.

Limitations in the scale and precision of current genome editing methods have motivated our recent work to develop a nuclease-independent eukaryotic multiplex genome engineering (eMAGE) platform^11^, which directs the annealing of multiple ∼90-nt single-stranded oligodeoxynucleotides (ssODNs) at the DNA replication fork to rapidly generate precise combinatorial (10^5^-10^6^) genome modifications with up to 40% efficiency across many (10-10^2^) loci in targeted genomic regions of *S. cerevisiae* (**Fig. 1a**). The eMAGE method uses a counter-selectable marker (*e.g*., *URA3*) adjacent to the targeted genomic region to enrich ssODN-edited cells via a co-selection strategy^11, 12^ (**Fig. 1b**). Like other ssODN-mediated genome editing methods^13–15^, eMAGE editing efficiency is greatly enhanced by inactivation of genes required for DNA mismatch repair (MMR)^11^. However, such constraints reduce genome stability, which could lead to the accumulation of unintended secondary mutations from unresolved DNA replication errors and spontaneous lesions, and requires prior modifications of the genome (*e.g*., deletion of MMR genes).

**Fig. 1:**
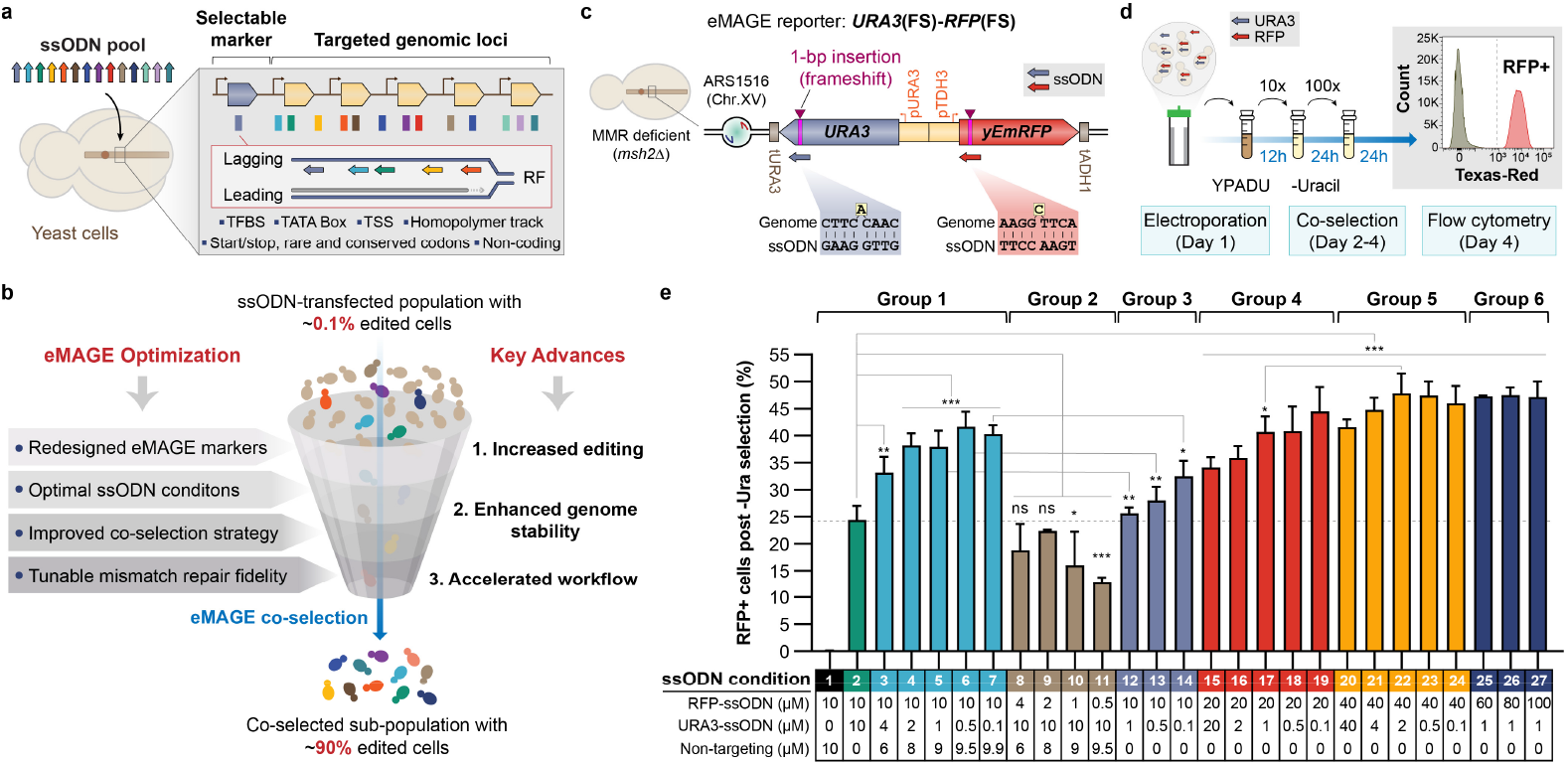
Increase eMAGE editing efficiency by redesigning co-selection markers and optimizing ssODN conditions. **a**, eMAGE efficiently generates precise combinatorial genome editing in diverse genetic elements through directing the simultaneous annealing of multiple transfected ssODNs to the lagging strand of a replication fork. eMAGE co-selection via an ssODN-edited adjacent selectable marker enriches cells with combinatorial editing in the targeted genomic loci. RF: replication fork. **b**, Systematic optimization of eMAGE at multiple levels by this study improves genome editing performance with faster editing workflow and enhances genome stability in eMAGE cells, resulting in ∼90% edited cells in the co-selected population. **c**, Fluorescent reporter design allows accurate and robust quantification of eMAGE editing efficiency. Yeast strain is deficient in MMR with deletion of the *MSH2* gene (*msh2*Δ). FS: frameshift. **d**, Typical eMAGE workflow for co-selection and flow cytometry of yeast strains carrying the *URA3*(FS)-*RFP*(FS) reporter, taking 3.5 days from ssODN electroporation to RFP readout. **e**, Comparison of eMAGE ARF across a series of conditions with different ssODN concentrations and ratios. All values represent mean ± SD for at least three replicates. p values of multiple-group comparisons from ordinary one-way ANOVA Dunnett’s test and p values of two-group comparisons from unpaired t-test. ns, not significant, *p < 0.05, **p < 0.01, ***p < 0.001.

In this study, we describe several key advances in eMAGE to address these limitations and enhance editing efficiency (**Fig. 1b**). First, we redesigned the eMAGE protocol and editing assay, substantially reducing experimental workflow time by two days, or 40%, and permitting rapid analysis of a larger number of edited cells with improved precision. Second, we performed systematic optimization of ssODN concentration, ratio and modification, as well as implemented a new co-selection strategy, leading to a striking increase in eMAGE efficiency with up to ∼90% edited cells in a population. Finally, we developed a system for transient and controllable MMR suppression by engineering expression of dominant negative MMR mutants, achieving high-efficiency editing in DNA repair-proficient cells with enhanced genome stability and reduced frequency of unintended secondary mutations.

We first set out to reduce co-selection time and increase assay throughput by developing an improved editing reporter compatible with flow cytometry to accelerate the eMAGE workflow. To measure eMAGE editing allelic replacement frequency (ARF), we previously used a *URA3-ADE2* reporter^11^ assay with a 5.5-day turnaround time to quantify *ADE2* ARF as the percentage of *ade2* edited cells (red colonies) on culture plates containing 5-fluoroorotic acid (5-FOA, a chemical compound that counterselects *ura3* cells). The redesigned eMAGE reporter harbors *URA3* and *yEmRFP* genes each carrying a predefined frameshift (FS) located in a previously characterized genomic locus^11^ of replication origin ARS1516 in chromosome XV (**Fig. 1c**). Yeast cells co-transfected with ssODNs restoring the correct reading frames of both *URA3* and *yEmRFP* were co-selected in liquid medium lacking uracil and analyzed via flow cytometry to determine the frequency of RFP positive cells, which represents eMAGE ARF (**Fig. 1d**). The new *URA3*(FS)-*RFP*(FS) reporter provides three advantages over the previous *URA3-ADE2* reporter^11^. First, it reduces culturing time needed for eMAGE coselection from four to two days by making use of the more robust positive *URA3* selection and rapid RFP readout (**Fig. 1d**), and avoiding laborious plating and colony screening procedures. Second, it eliminates interference from secondary mutations due to the extremely low frequency of spontaneous frameshift correction (which we did not observe), rendering co-selection more stringent by avoiding background 5-FOA resistant mutants (∼10% of total *ura3* colonies^11^) that are unrelated to ssODN editing. Third, it facilitates rapid (∼30 seconds per sample) and more accurate quantification of eMAGE editing frequency via analyzing a larger population by flow cytometry (>10^5^ flowed cells vs. 10^2^-10^3^ colonies), resulting in a significantly improved assay throughput (>100 samples per hour). Together, this fluorescent reporter system permits convenient, fast and accurate comparison of eMAGE ARF among diverse conditions, establishing a facile system for optimization.

Using the fluorescent reporter system, we next attempted to optimize eMAGE ARF by investigating a wide range of concentrations between ssODNs targeting the edited locus (*RFP*) and the co-selection marker (*URA3*). Holding concentration of the RFP-ssODN at 10 μM and total ssODN concentration at 20 μM by supplementing a non-targeting ssODN, we decreased concentration of the URA3-ssODN and observed up to ∼40% ARF in the *RFP* gene. This is a significant increase over the 25% baseline using the previous protocol^11^, in which ssODN concentration is 10 μM for both the coselection marker (*URA3*) and editing targeted site (*RFP*) (**Fig. 1e**, group 1). Conversely, samples with a decreased RFP-to URA3-ssODN concentration ratio showed reduced eMAGE ARF in the range of ∼13-22% (**Fig. 1e**, group 2). These results suggest that eMAGE ARF can be enhanced by limiting the cellular availability of ssODN targeting the selectable marker to increase the stringency of co-selection. With the same ratio of ssODN concentrations, total concentration of ssODNs through 40 μM positively correlated with eMAGE ARF up to ∼48% (**Fig. 1e**, compare groups 1, 3, 4 and 5). Further increase of RFP ssODN concentration to 60, 80 or 100 μM showed no further enhancement in eMAGE ARF (**Fig. 1e**, group 6), despite a slight increase in the RFP positive frequency in ssODN-transfected cells prior to *URA3* coselection (**Extended Data Fig. 1a**), suggesting that eMAGE ARF reaches an efficiency ceiling with URA3-ssODN at ∼40 μM. Given that concentration of URA3-ssODN also positively influences the initial number of URA3 cells (**Extended Data Fig. 1b**) and the total ssODN concentration negatively correlates with cell survival after electroporation (**Extended Data Fig. 1c**), we converged on optimal ssODN conditions: 20-60 μM total ssODN with a 20:1 ratio of ssODNs targeting edited site(s) and the co-selection marker (*e.g*., **Fig. 1e**, condition 22). We also attempted to increase the nuclease resistance of ssODNs by modifying their terminal nucleotides at both ends with phosphorothioate bonds and observed a minor but statistically significant ARF increase of ∼5% over unmodified ssODNs at 40 and 80 μM (**Extended Data Fig. 1d**).

We previously observed that eMAGE ARF decreases as the distance between edited sites and the co-selection marker increases: ∼40% at 1 kb, to ∼15% at 5 kb, to ∼ 5% at 20 kb^11^. We hypothesize that flanking the targeted genomic region with two co-selection markers would sustain high levels of multisite editing across a longer genomic locus. To test this hypothesis, we constructed a five-gene construct *URA3*(FS)-*BGR*(FS)-*ADE2*(FS), comprising three fluorescent reporters (*ymTagBFP2, ymUkG1* and *yEmRFP*) flanked by two selectable markers (*URA3* and *ADE2*), each of which bears a predefined frameshift mutation (**Fig. 2a**). Applying the ssODN concentration and ratio from the optimization experiments described above (1 μM for each co-selection marker and 20 μM for each fluorescent reporter), we observed that co-selection using both *URA3* and *ADE2* markers led to significantly higher eMAGE ARF (50-70%) in all three fluorescent reporters, as compared to coselection using a single marker (**Fig. 2b**). Analysis of eMAGE multiplex editing further revealed that dual-marker co-selection enriched significantly more double- and triple-edited cells (∼60% vs. ∼30-40%) and dramatically reduced the unedited sub-population from ∼30% to ∼5% (**Fig. 2c**). Using the three fluorescent genes in the eMAGE reporter, we further compared three genetic architectures for dual-marker co-selection by differentially gating the flow cytometry data to determine how the relative positions of the co-selection markers affect ARF (**Extended Data Fig. 2a**). We found that dual-marker co-selection in any placement configuration outperformed single-marker co-selection, and that markers flanking the targeted site resulted in the highest ARF (**Extended Data Fig. 2b**).

**Fig. 2:**
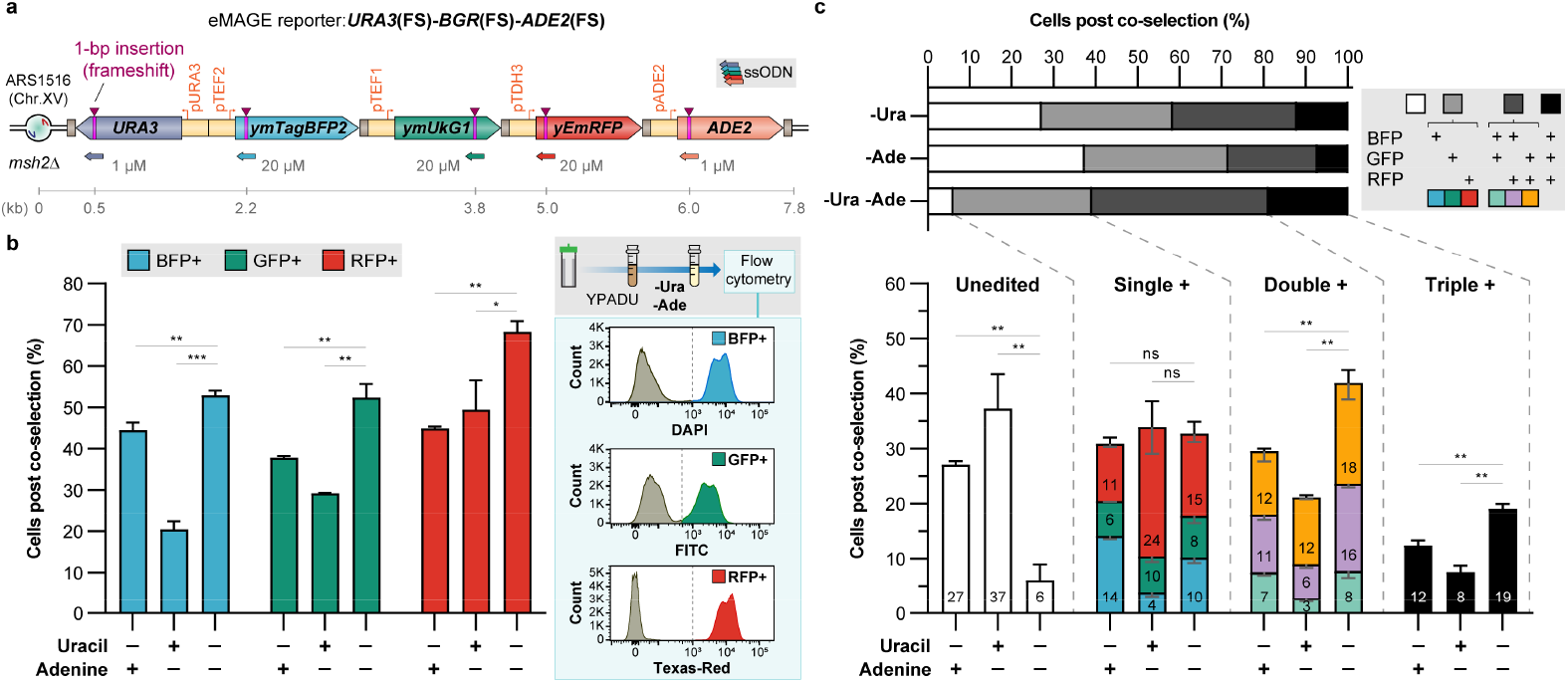
Improved eMAGE multiplex editing with dual selectable markers. **a**, eMAGE reporter carrying three fluorescent and two selectable marker gene cassettes for rapid quantification of eMAGE multiplex ARF by flow cytometry. Sizes of the genetic elements are not shown in actual scale. Distance of frameshift mutations in each targeted gene to the replication origin is shown. **b**, Comparison of eMAGE ARF of three fluorescent genes in cells co-selected using either single or dual selectable markers. Typical eMAGE workflow and representative flow cytometry plots of cells passed through -Ura -Ade co-selection are shown on the right. **c**, Multiplex editing analysis of cells carrying the eMAGE reporter shown in **a**. Co-selected cells have different combinations of fluorescent signals based on the frameshift correction status in three fluorescent reporter genes. The internal numbers of the lower stack bars represent mean percentages of the sub-populations with different fluorescent signal combinations. The internal numbers of the lower stack bars represent mean percentages of the sub-populations with different fluorescent signal combinations. The upward facing error bars on top of the stacked bars represent SD of the mean percentages of the four editing categories (unedited, single-, double-, triple-edited cells). The downward facing error bars internal to the stacked bars represent SD of the mean percentages of each sub-population with the specified multiplex editing statuses by the color codes. All values represent mean ± SD for at least three replicates. Statistical significance analysis of the means of each editing category was performed. p values from ordinary oneway ANOVA Dunnett’s test. ns, not significant, *p < 0.05, **p < 0.01, ***p < 0.001.

Consistent with prior observations^11, 13^, we confirmed that deletion of *MSH2* (*msh2*Δ) increased editing frequency three-to five-fold in all five markers of the multi-gene *URA3*(FS)-*BGR*(FS)-*ADE2*(FS) reporter (**Fig. 2a**) after transfection of a pool of five ssODN at the optimal concentrations (*i.e*., 1 μM for *URA3, ADE2* and 20 μM for *B*-, *G*-, *RFP* genes) (**Extended Data Fig. 3**). This *msh2*Δ background also led to a 10-fold increase of double-edited URA3, ADE2 cells, bolstering the dual-marker co-selection strategy. To maintain high eMAGE ARF (∼90% edited) while circumventing permanent MMR inactivation, we adopted a similar approach used successfully for MAGE in bacteria^16^: transient expression of dominant negative mutants of MMR proteins (MMR-DN) in DNA repair-proficient cells to temporarily suppress MMR prior to ssODN electroporation (**Fig. 3a**). To gain tighter control of the expression level of MMR-DN mutants and enhance the reproducibility of this method, we modified a previously reported β-estradiol inducible expression system by stably integrating the estradiol-responsive transcription activator (GEM) expression cassette in a defined genomic locus (YFL033C^18^) to constitutively express GEM at a moderate level in order to achieve tunable induction of MMR-DN upon β-estradiol titrations. We then empirically characterized this regulator using RFP expression which showed a dynamic range of inducible expression levels with no detectable uninduced basal expression. (**Extended Data Fig. 4a**).

**Fig. 3:**
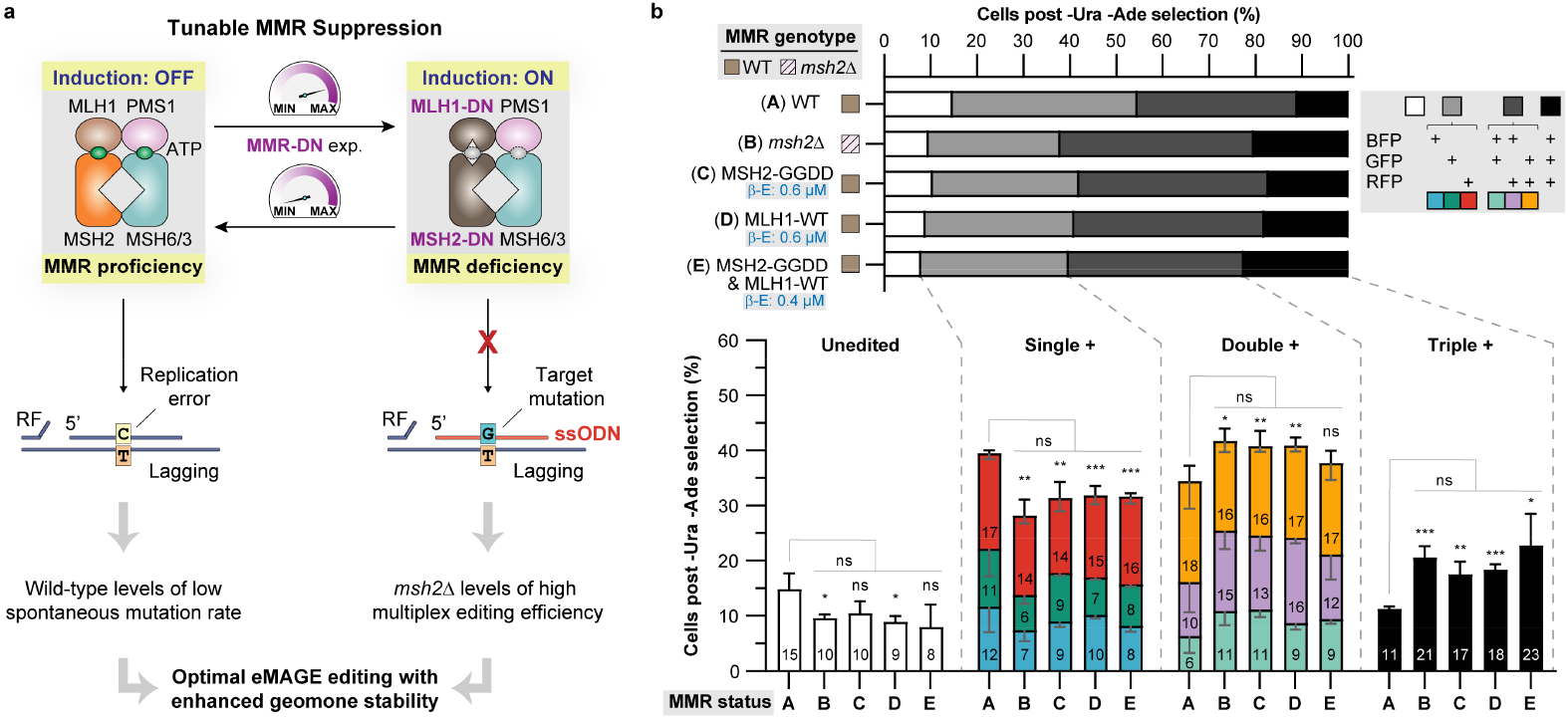
Transient inactivation of mismatch repair enables efficient eMAGE editing in DNA repair-proficient cells. **a**, Rationally designed tunable overexpression of dominant negative subunits of the yeast DNA mismatch recognition and repair complex MutSα/β causes transient MMR deficiency in DNA repair-proficient cells, resulting in high on-target editing efficiency with low spontaneous mutation rate. MMR: DNA mismatch repair. DN: dominant negative. RF: replication fork. **b**, Multiplex editing analysis of cells expressing selective MMR-DN mutants at three selective conditions. β-E: β-estradiol. The internal numbers of the lower stack bars represent mean percentages of the sub-populations with different fluorescent signal combinations. The internal numbers of the lower stack bars represent mean percentages of the sub-populations with different fluorescent signal combinations. The upward facing error bars on top of the stacked bars represent SD of the mean percentages of the four editing categories (unedited, single-, double-, triple-edited cells). The downward facing error bars internal to the stacked bars represent SD of the mean percentages of each sub-population with the specified multiplex editing statuses by the color codes. All values represent mean ± SD for at least three replicates. Statistical significance analysis of the means of each editing category was performed. p values from ordinary one-way ANOVA Dunnett’s test. ns, not significant, *p < 0.05, **p < 0.01, ***p < 0.001.

Next, we screened nine different MMR-DN mutants from three subunits (MSH2, MSH6 and MLH1) of the yeast mismatch repair MutS complexes, which have shown strong dominant negative effects in previous studies^19–21^, together with three wild-type subunit proteins (**Extended Data Table 1**),via episomal overexpression in a MMR-proficient strain (**Extended Data Fig. 4b**) harboring the *URA3*(FS)-*BGR*(FS)-*ADE2*(FS) reporter (**Fig. 2a**). Among the 12 tested MMR protein variants, moderate expression (β-estradiol: 0.4 μM) of MSH2-DN mutants^19^ led to the highest frameshift correction frequencies in three fluorescent genes, followed by MLH1-WT (for which overexpression was reported to have a DN effect^20^) and MLH1-DN mutants^20^, while expression of MSH6-DN mutants^21^ surprisingly showed no significant dominant negative effect in our assay (**Extended Data Fig. 4c**). Notably, we screened MMR-DN candidates using the frequency of fluorescent cells in ssODN-transfected population prior to co-selection, which directly reflects the efficiency of ssODN incorporation and is thus more sensitive to MMR suppression than the post co-selection eMAGE ARF.

We then selected MSH2-GGDD and MLH1-WT for further optimization by fine-tuning the expression level of both mutants alone and in combination. Expression of MSH2-GGDD alone at relatively high levels (β-estradiol: 0.6-1 μM) or in combination with MLH1-WT at relatively low levels (β-estradiol: 0.2-0.4 μM) yielded similar frameshift correction frequency as the *msh2*Δ cells with no statistically significant difference (**Extended Data Fig. 4d**). Importantly, analysis of eMAGE multiplex editing demonstrated that three selected MMR-DN mutant expression conditions each confer proportions of unedited, single-, double-, and triple-positive cells indistinguishable from *msh2*Δ background (**Fig. 3b**). Interestingly, albeit having significantly less total edited cells than other strains with permanent or transient MMR inactivation, the MMR-proficient (WT) strain is capable of considerably improved eMAGE editing as compared to our prior results^11^, due to the optimization of ssODN condition and the new eMAGE co-selection design described in this study (**Fig. 1** and **2**). This provides a new opportunity to conveniently apply the reported optimized conditions to enable eMAGE with modest efficiency in DNA repair-proficient cells independent of MMR modulation. Assessment of spontaneous mutation rate by Luria-Delbrück fluctuation analysis^22^ revealed that MMR-proficient cells carrying episomal MSH2-GGDD have a similar frequency of spontaneous mutations to wild-type cells, which can be transiently toggled to a level similar to *msh2*Δ cells upon MMR-DN expression (**Extended Data Table 2**, column A). Further analysis of data collected in this study demonstrates that tunable expression of MSH2-GGDD not only preserves wild-type levels of low spontaneous mutations (**Extended Data Table 2**, column B), but also inducibly enables *msh2*Δ levels of high-efficiency multiplex editing in a transiently relaxed genomic context with up to 10^4^ possible combinations of genomic diversity (**Extended Data Table 2**, column C-E).

In summary, this study accelerates the eMAGE editing workflow by ∼40%, improves on-target editing efficiency up to 70% for single edits and >60% for multiplex editing, resulting in ∼90% of cells carrying intended genomic modifications after one round of eMAGE. The engineered MMR system for transient suppression of DNA repair genes permits high-efficiency eMAGE editing in DNA repair-proficient cells, reducing unintended mutations by at least 4-fold during the editing process and ∼17-fold in long-term outgrowth, thereby stabilizing the genome in eMAGE strains (**Fig. 1b**). We expect that these advances will empower eMAGE for tasks that demand precise and combinatorial diversification of targeted genetic elements with prolonged cultivation^23^, such as protein engineering, molecular evolution, heterologous pathway optimization, and synthetic genome construction. Given the highly conserved nature of eukaryotic DNA replication and repair, we envision that the reported advances of eMAGE can guide subsequent development of analogous nuclease-independent genome editing technologies in other yeast and eukaryotic cells.

## Methods

### Strain construction

Yeast strains used in this study can be found in **Supplementary Table 1** and sequences of important genomic loci can be found in **Supplementary File 1**. To construct the yeast strain harboring the *URA3*(FS)-*RFP*(FS) eMAGE reporter (**Fig. 1c**), the *URA3* cassette of EMB101^11^ was first replaced by a PCR product of *ymUkG1*^24^(FS)-*yEmRFP*^25^(FS) cassettes via standard homologous recombination (HR) and 5-FOA counterselection, resulting in yeast strain SZL149. Next a *URA3* cassette was reintroduced to replace the *ymUKG1*(FS) cassette via HR and uracil drop-out selection, resulting in strain SZL238 with *URA3-yEmRFP*(FS). Next a frameshift mutation was introduced in the *URA3* coding sequence by eMAGE with an ssODN, OZL327 (information of PCR primers can be found in **Supplementary Table 3**), followed by 5-FOA counterselection, resulting in strain SZL247. Finally, the *MSH2* gene was deleted by a *KanMX4* cassette conferring G418 (geneticin) resistance, resulting in strain SZL335 that was used in **Fig. 1e**. To construct strains capable of β-estradiol inducible expression, GEM effector expression cassette together with the *LEU2* marker was PCR amplified from pHES839^17^ (Addgene # 87941) using primers OZL348 and OZL349 and integrated downstream of the *YFL033C* locus of SZL149 and SZL281 (SZL149+*msh2*Δ), resulting in strain SZL281 and SZL308, respectively. To construct the yeast strains harboring the *URA3*(FS)-*BGR*(FS)-*ADE2*(FS) eMAGE reporter (**Fig. 2a**), *URA3-ymTagBFP2*(FS) cassettes (*ymTagBFP2*: codon-optimized *mTagBFP2*^26^ for *S*. *cerevisiae* from this study, coding sequence can be found in **Supplementary File 1**) was integrated between ARS1516 and the *ymUkG1*(FS) cassette of SZL281 and SZL308, resulting in strain SZL345 and SZL347, respectively. Next a frameshift mutation was introduced in the *URA3* and *ADE2* coding sequences by a round of eMAGE with ssODN OZL327 and OZL428, resulting in strain SZL376 that was used in **Fig. 3b** and **Extended Data Fig. 3c,d** (MMR-WT), and strain SZL348 that was used in **Fig. 2b,c**, **Fig. 3b** and **Extended Data Fig. 3c,d** (*msh2*Δ). Yeast transformation was performed via the optimized PEG/lithium acetate protocol of the Wilson lab^27^. All eMAGE reporter loci were sequence verified in the final strains.

### Media

For yeast cultivation, cells were grown in nonselective YPADU liquid medium, which consists of YPD (10 g/l Yeast Extract, 20 g/l Peptone, 20 g/l Dextrose), supplemented with 40 mg/l adenine hemi sulfate, and 20 mg/l uracil. For auxotrophic selection, synthetic defined media with 2% glucose and drop-out of corresponding nutrients were used. For eMAGE with frameshifted markers, yeast was cultured in YPADU and recovered immediately after ssODN electroporation in YPADU with 0.5 M sorbitol. For culturing yeast strains carrying episomal expression vectors of MMR genes, YPADU with 200 μg/ml zeocin was used.

### Plasmid cloning

Plasmids used in this study can be found in **Supplementary Table 2**. For inducible expression of MMR subunits, the *LEU2* cassette of pRSII425^28^ (Addgene #35468) was replaced by a Sh *ble* cassette conferring zeocin resistance, resulting in a shuttle vector pRSII42B (this study) harbored in strain SZL134. Wild-type MMR genes (*MSH2, MSH6, MLH1*) were PCR amplified from genomic DNA of yeast strain BY4741^29^ and cloned together with the bidirectional inducible *Gal1-10* promoter into the PCR amplified pRSII42B backbone with OZL14 and OZL15 via Gibson assembly^30^ using NEBuilder Kit (NEB, catalog E2621). MMR-DN mutant genes were subsequently generated via site-directed mutagenesis using either Q5 Site-Directed Mutagenesis Kit (NEB, catalog E0554S) or QuikChange Lightning Multi Site-Directed Mutagenesis Kit (Agilent, catalog 210515) according to the manufacturer protocols.

### ssODN electroporation

ssODN electroporation was carried out using the previously reported protocol^11^. In brief, a single colony was inoculated in a 2 ml starter culture of appropriate media and grown overnight to saturation. The next day, the culture was diluted 1:100 into 10 ml fresh media and grown until the OD600 is between 0.6 and 1.0 in 4-6 hours. Cells were then pelleted and incubated in DTT-LiTE buffer (100 mM lithium acetate, 25 mM DTT, 500 mM hydroxyurea, 1x TE at pH 8.0) for 30 min at 30°C. Cells were then washed with ice cold water and 1 M sorbitol. The specified combinations and concentrations of ssODNs were prepared in 200 ul 1 M sorbitol, and used to resuspend the washed cells. ssODN electroporation was carried out in a 2-mm cuvette (Biorad, catalog 1652086) at 1500 V, 25 μF, 200 Ω. Immediately after electroporation, 1 ml eMAGE recovery medium (YPADU with 0.5 M sorbitol) was added to the cuvette, and the resuspended cells were added to a 15 ml culture tube containing 4 ml recovery media (final volume: 5 ml). Cells were then incubated at 30°C in a drum rotator (Fisher Scientific, catalog 14251-228Q/ 232Q) for 12 hours prior to co-selection. Information of ssODNs used in this study can be found in **Supplementary Table 3**.

### eMAGE co-selection

After overnight recovery after ssODN electroporation, 500 μl of saturated culture was spun down, washed once with 1 ml sterile water, and transferred to 5 ml synthetic defined media with drop-out of auxotrophic nutrients based on the applicable selectable markers. This 10-fold diluted culture was first grown for 24 hours (single-marker co-selection) or 48 hours (dual-marker co-selection), then diluted 100-fold in 3 ml of the same synthetic defined media and incubated for another 24 hours to saturation. Sequential dilutions of 10- and 100-fold in two steps ensure that unedited cells represent less than 0.1% in co-selected population and therefore exert negligible influence on eMAGE ARF quantification, while making sure sampling errors in each dilution are within acceptable range (95% confidence level and 2% margin of error).

### Flow cytometry analysis

Flow cytometry was carried out on BD FACSAria II. ymTagBFP2 was excited by the violet laser (405-nm) with the DAPI emission filter (450/40); ymUkG1 was excited by the blue laser (488-nm) with the FITC emission filter (530/30); yEmRFP was excited by the green laser (561-nm) with the Texas-Red emission filter (610/20). Yeast cells were washed and resuspended in 1x PBS prior to flow cytometry. Flow cytometry data were analyzed in FlowJo V10.

### Cell survival assay

To measure the effect of ssODN concentration on cell survival, strain SZL335 was prepared the same way as in eMAGE and electroporated with representative concentrations of RFP- and URA3-ssODN. Immediately after ssODN electroporation, cells were plated onto YPADU plate with 0.5 M sorbitol and incubated at 30°C for 2 days. The number of colonies on each plate was counted and normalized with the plating volumes and dilution factors to calculate the percentage of viable cells in each condition. Cell viability data were normalized to untreated cells (100 % viable).

### β-estradiol inducible expression

The β-estradiol inducible system was constructed as above (Strain construction and Plasmid cloning). Constitutively expressed GEM translocates from cytosol to nucleus upon β-estradiol binding and activates transcription of the *Gal1-10* promoter. β-estradiol (Sigma, catalog E1024) was prepared as a 10 mM stock solution in 100% ethanol, then diluted in media to working concentrations between 0 and 1 μM. Titration of inducible expression levels was empirically determined using RFP expression. Plasmids were maintained via zeocin selection. RFP fluorescence and cell density (OD600) were measured in each β-estradiol concentration along a 24-hour time course using a BioTek plate reader (Synergy H1), and selective time points were plotted to visualize the dose-response curve of gene expression as shown in **Extended Data Fig. 4a**.

### Fluctuation analysis

To quantify the spontaneous mutation rate of selective eMAGE strains in **Fig. 3d**, Luria-Delbrück fluctuation analysis was performed using a protocol modified from Lang^22^. In brief, a 2 ml starter culture was inoculated from a single colony in synthetic defined medium with drop-out of uracil and grown overnight to saturation. The next day different cultures were adjusted to the same OD of 1.0 and then diluted 5,000-fold in nonselective YPADU medium without or with β-estradiol in the specified concentration. 100 ul of diluted yeast culture was added to each well of a 96-well plate at a seeding density of 200-600 cells and incubated at 30°C without shaking for 2 days until saturation. For each 96-well plate, cultures of 12 wells were pooled and used to determine the average number of cells per culture (**N**) by limited-dilution plating. The entire cultures (100 μl) of the remaining 84 wells were spot plated on 5-FOA media and incubated at 30°C for 4 days. Mutation rate (**μ**): mutation events per cell per division or generation, was calculated using the *p0* method with formula: μ = −ln(*p*0)/N, where ***p*0** is the fraction of spot cultures with no colony.

### Calculations of spontaneous mutation and eMAGE on-target editing

In **Extended Data Table 2**, spontaneous mutations rates of the *URA3* gene shown in column A were determined by the fluctuation analysis described above. Estimated numbers of spontaneous mutations shown in column B were calculated by normalizing the size of the *URA3* gene (804 bp) to S288c genome (12 mb) and multiplying the numbers of cell division in a typical round of eMAGE (∼66 cell divisions) or after accumulative cultivation of 10 days (∼120 cell divisions). The cell division numbers were estimated based on the total time of yeast cultivation (*i.e*., ∼132 hours during eMAGE comprising 48-hour: isolated colony forming from frozen glycerol stock, 24-hour: eMAGE starter culture incubation, 12-hour: recovery post ssODN electroporation, and 48-hour: eMAGE co-selection) with an average 2-hour doubling time. The Average numbers of eMAGE on-target edits shown in column C were derived from data of the eMAGE multiplex editing experiments shown in **Fig. 3b** using formula: Avg. edits= Σ[(% of sub-population)×(number of edits)], where ‘% of sub-population’ is the percentage of cells with either 0, 1, 2 or 3 edits. The number of double-edited URA3, ADE2 cells after ssODN electroporation and recovery shown in column D were calculated with cell viability data (∼10% at 60 μM ssODN) shown in **Extended Data Fig. 1c** and the *URA3, ADE2* double frameshift correction frequency (∼0.0015% for WT, ∼0.015% for *msh2*Δ and MSH2-GGDD (β-E: 0.6 μM)) shown in **Extended Data Fig. 3**. Total numbers of eMAGE edited sites shown in column F were calculated by multiplying the values in column C and D.

### Statistical analysis

GraphPad Prism 8 was used for all statistical analysis. All data sets were generated with at least three replicates unless specified otherwise, and error bars are reported as mean ± SD. Ordinary one-way ANOVA Dunnett’s tests were used to measure significance of data of multiple groups, and unpaired t-tests were used to measure significance of data of two groups with confidence level cutoff of ns, not significant ≥ 0.05, *p < 0.05 **p < 0.01, ***p < 0.001, ****p < 0.0001.

## Acknowledgements

We thank Dr. Edward Barbieri for sharing his collection of yeast strains and prior experience in eMAGE protocol optimization; Kenneth Nelson (Yale West Campus Analytical Core facility) for assistance with flow-cytometry experiments; Dr. Neta Dean (Stony Brook University) for sharing the yEmRFP construct. We are grateful to members of the Isaacs and Rinehart laboratories and Brett Lindenbach for their critical discussions and feedback; members of MacMicking, Chen and Levchenko laboratories for their advices and shared reagents for experiments; and members of the Lindenbach laboratory for reviewing the manuscript. We gratefully acknowledge support from the NSF (IOS-1923321) and Emerson Collective for funding.

## Author Contributions

F.J.I and Z.L. conceived the experiments and directed the studies. Z.L. and E.M. conducted the experiments and analyzed data. Z.L., E.M. and F.J.I wrote the manuscript.

## Data availability

The plasmids generated from this study will be deposited on Addgene (http://www.addgene.org) upon publication. Yeast strains used in this study are available upon request to the corresponding authors.

---

**Extended Data Table 1:** Subunit variants of DNA mismatch repair MutS complexes tested in this study.

| MMR gene | Variant | Mutated domain | Protein biochemistry | Reported spontaneous mutation rate<sup>a</sup> (vs. wild-type) | Sources |
| --- | --- | --- | --- | --- | --- |
| MSH2 | Wild-type | - | - | Unchanged<sup>b</sup> | Alani et al, *Mol Cell Biol.* 1997<sup>19</sup> |
|  | G693D | ATPase (MSH2) | Defective ATP binding | ~50-fold<sup>b</sup> |  |
|  | G855D |  | Defective ATP binding, hydrolysis and MutS-DNA complex formation |  |  |
|  | G693D, G855D (GGDD) |  | Combined effects of two single mutants |  |  |
| MSH6 | Wild-type | - | - | Unchanged<sup>b</sup> | Gupta et al, *Nat Genet.* 2000<sup>21</sup> |
|  | S1036P | ATPase (MSH6) | Defective ATP hydrolysis | ~15-fold<sup>b</sup> |  |
|  | G1067D | Domain interacting with ATPase (MSH2) | Defective ATP hydrolysis and MutS-DNA complex formation |  |  |
|  | G1142D |  |  |  |  |
| MLH1 | Wild-type | - | Imbalanced MLH1-PSM1 stoichiometry | ~13-fold<sup>b</sup> | Shcherbakova et al, *Mol Cell Biol.* 1999<sup>20</sup> |
|  | G64R | ATPase (MLH1) | Defective ATP binding and hydrolysis | ~6-fold<sup>c</sup> |  |
|  | I65N |  |  |  |  |
|  | T114M |  |  |  |  |

<sup>a</sup> Based on reported data of the referenced literatures, showing the fold increases of mutation rate upon overexpression of specific MMR proteins as compared to the corresponding wild-type strains.
<sup>b</sup> Overexpressed from high-copy 2 $\mu$ plasmids. Mutation rates of *CAN1* gene were measured by quantification of canavanine resistance colonies.
<sup>c</sup> Expressed from the endogenous *MLH1* locus. Mutation rates of *lys2::InsE-A14* locus were measured by quantification of Lys<sup>+</sup> colonies.

**Extended Data Table 2:**
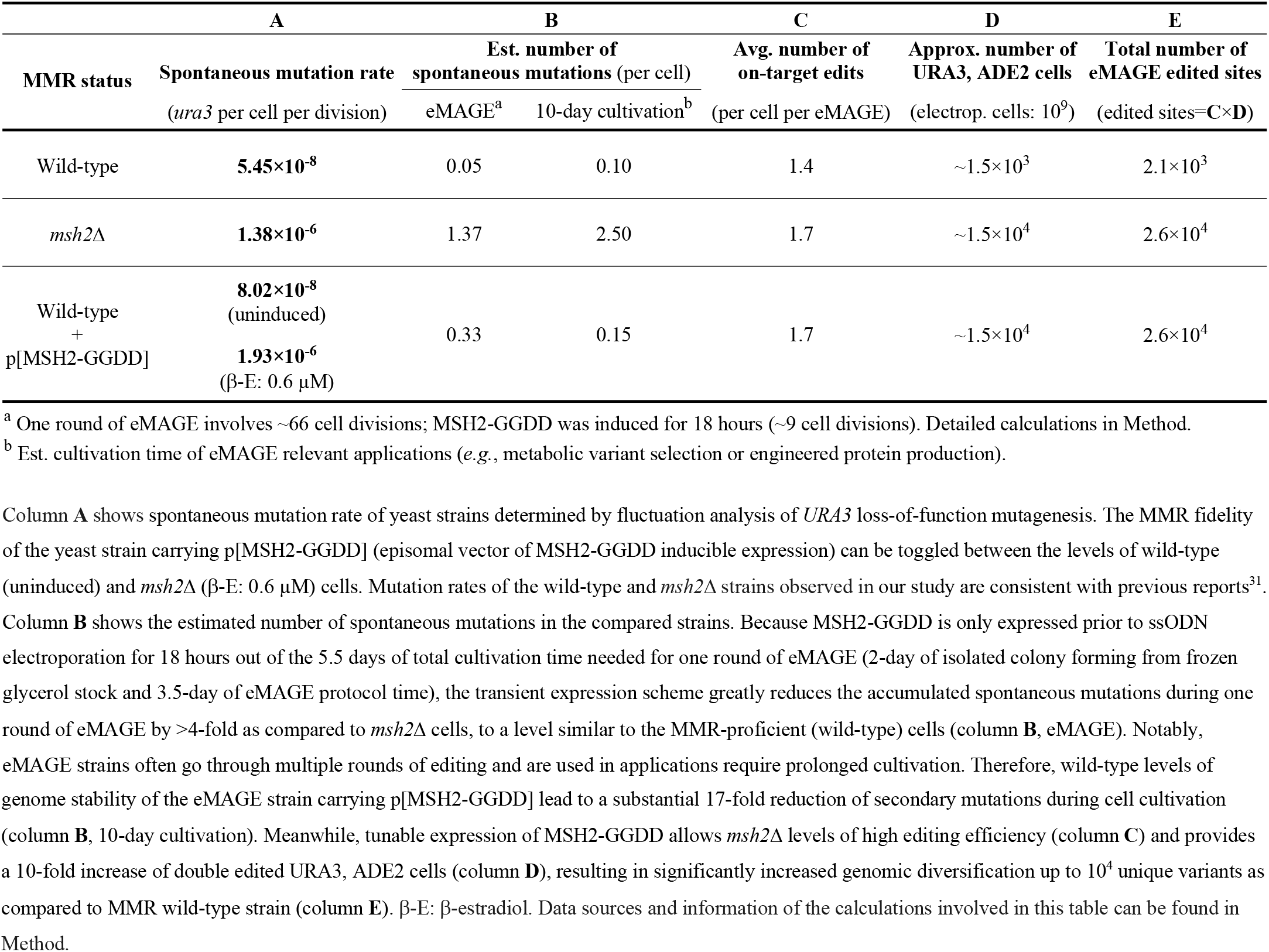
Influence of MMR proficiency on eMAGE genome editing and spontaneous mutations.

**Extended Data Fig. 1:**
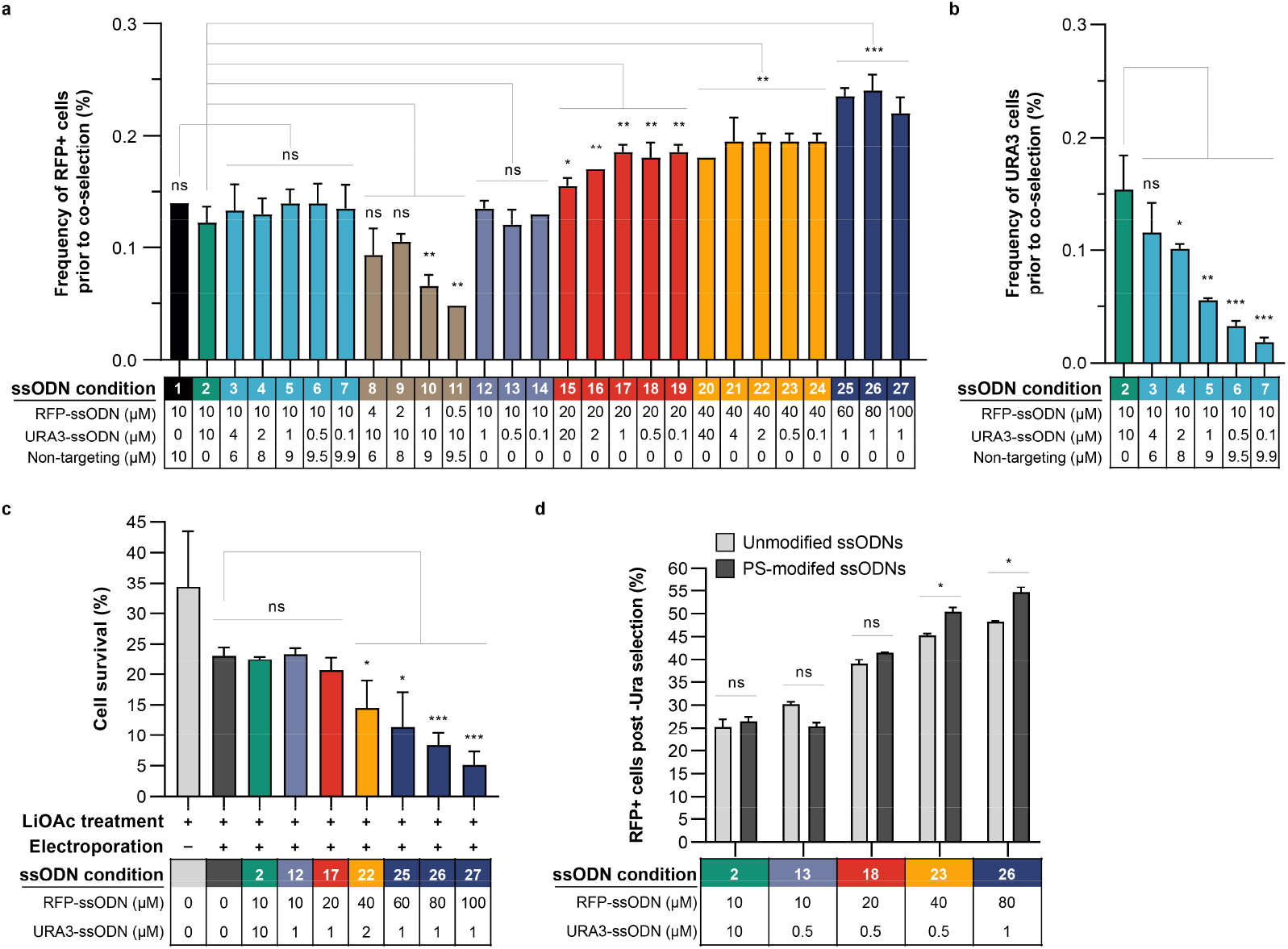
Effect of ssODN concentration and modification on frequency of edited cells and cell viability. **a**, Frequency of RFP positive cells in ssODN-transfected population prior to -Ura selection under different ssODN concentrations and ratios used in **Fig. 1e**. It positively correlates with the concentration of RFP-ssODN. **b**, Frequency of URA3 cells decreases with URA3-ssODN concentration. **c**, Cell survival of selective conditions used in **Fig. 1e**, showing decreased viability after electroporation with increased ssODN concentration. Cell viability data were normalized to untreated cells (100 % viable). **d**, Protecting ssODN from exonuclease degradation by modifying the last 4 nt of its 5’ and 3’ ends with phosphorothioate bonds does not result in major enhancement of eMAGE ARF as compared to unprotected ssODNs. All values represent mean ± SD for at least three replicates. p values of multiple-group comparisons from ordinary one-way ANOVA Dunnett’s test and p values of two-group comparisons from unpaired t-test. ns, not significant, *p < 0.05, **p < 0.01, ***p < 0.001.

**Extended Data Fig. 2:**
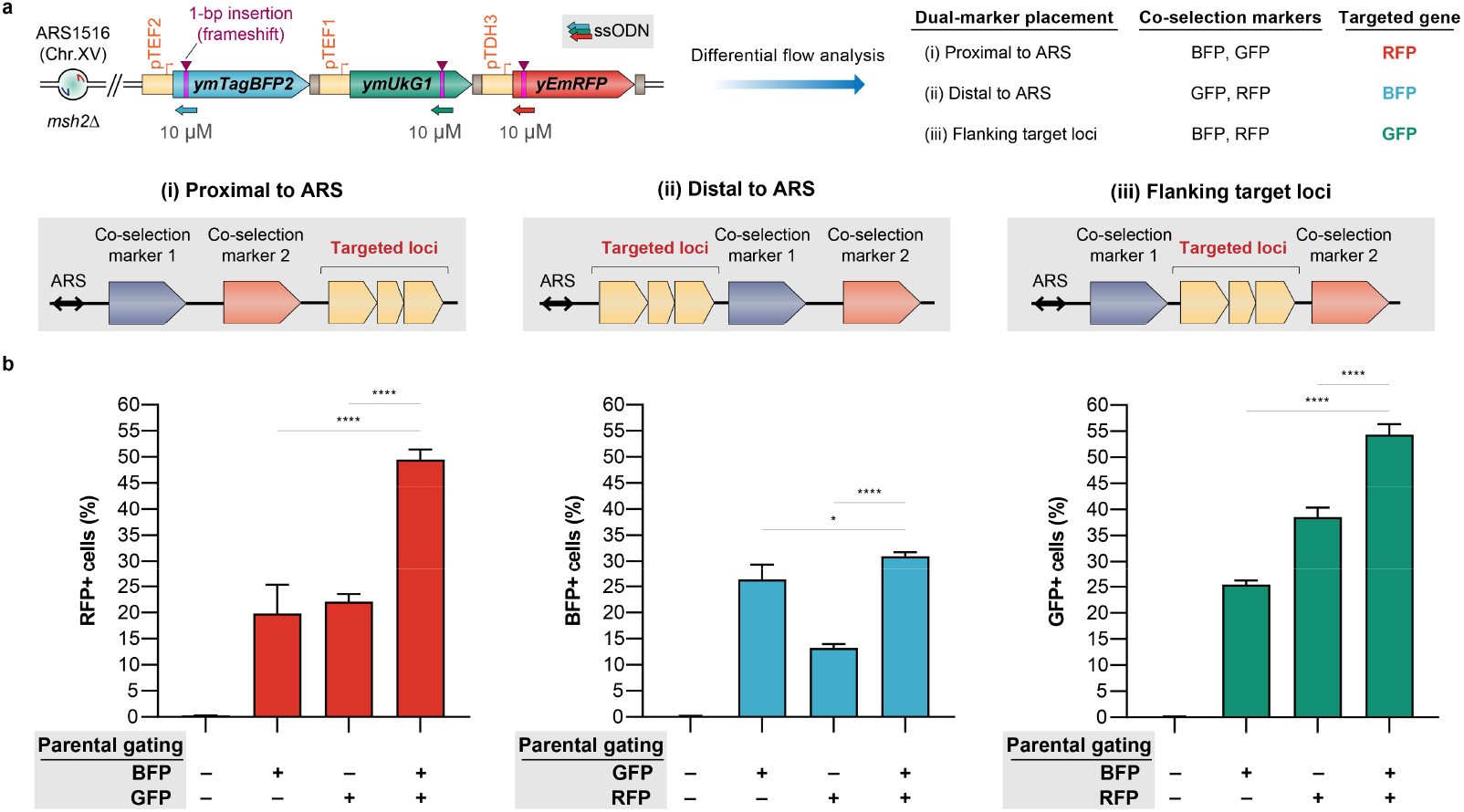
Comparison of three dual-marker placement strategies. **a**, eMAGE reporter used in this analysis containing the same frameshifted fluorescent reporter genes as the one in **Fig. 2a**. To simulate the three possible architectures for co-selection marker placement: (i) proximal to ARS, (ii) distal to ARS, and (iii) flanking targeted loci, the flow cytometry data collected from the same experiments were analyzed as follows: two of the three fluorescent markers were used as parental gates to select the cells with either single or double positive fluorescent phenotypes. The percentage of these gated cells that are positive for the remaining (third) fluorescent gene represents the eMAGE coselection ARF. **b**, Comparison of eMAGE ARF using co-selection with either a single or dual fluorescent marker in the above three placement configurations. In all scenarios, co-selection with two markers (last bar) yields higher eMAGE ARF compared to co-selection with only a single marker. Architecture (iii), in which the co-selection markers flank the targeted loci, resulted in the highest ARF gain among the tested dual-marker placement strategies. All values represent mean ± SD for at least three replicates. p values from ordinary one-way ANOVA Dunnett’s test. *p < 0.05, **p < 0.01, ***p < 0.001, ****p < 0.0001.

**Extended Data Fig. 3:**
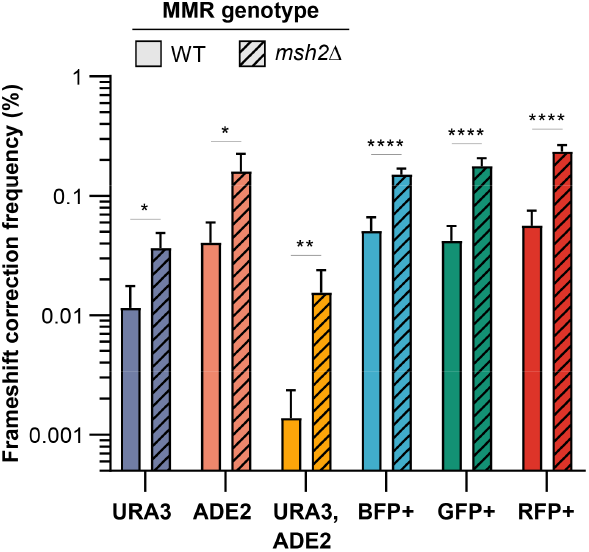
MMR deficiency promotes eMAGE editing and benefits implementation of dual-marker co-selection. **a**, Comparison of MMR wild-type (WT) and *MSH2* knockout (*msh2*Δ) strains for their frameshift correction frequency of each marker in the eMAGE reporter shown in **Fig. 2a**. Deletion of *MSH2* confers a three to five-fold increase in frameshift correction frequency and a 10-fold increase of double-edited URA3, ADE2 cells. All values represent mean ± SD for at least three replicates. p values from unpaired t-test. *p < 0.05, **p < 0.01, ***p < 0.001, ****p < 0.0001.

**Extended Data Fig. 4:**
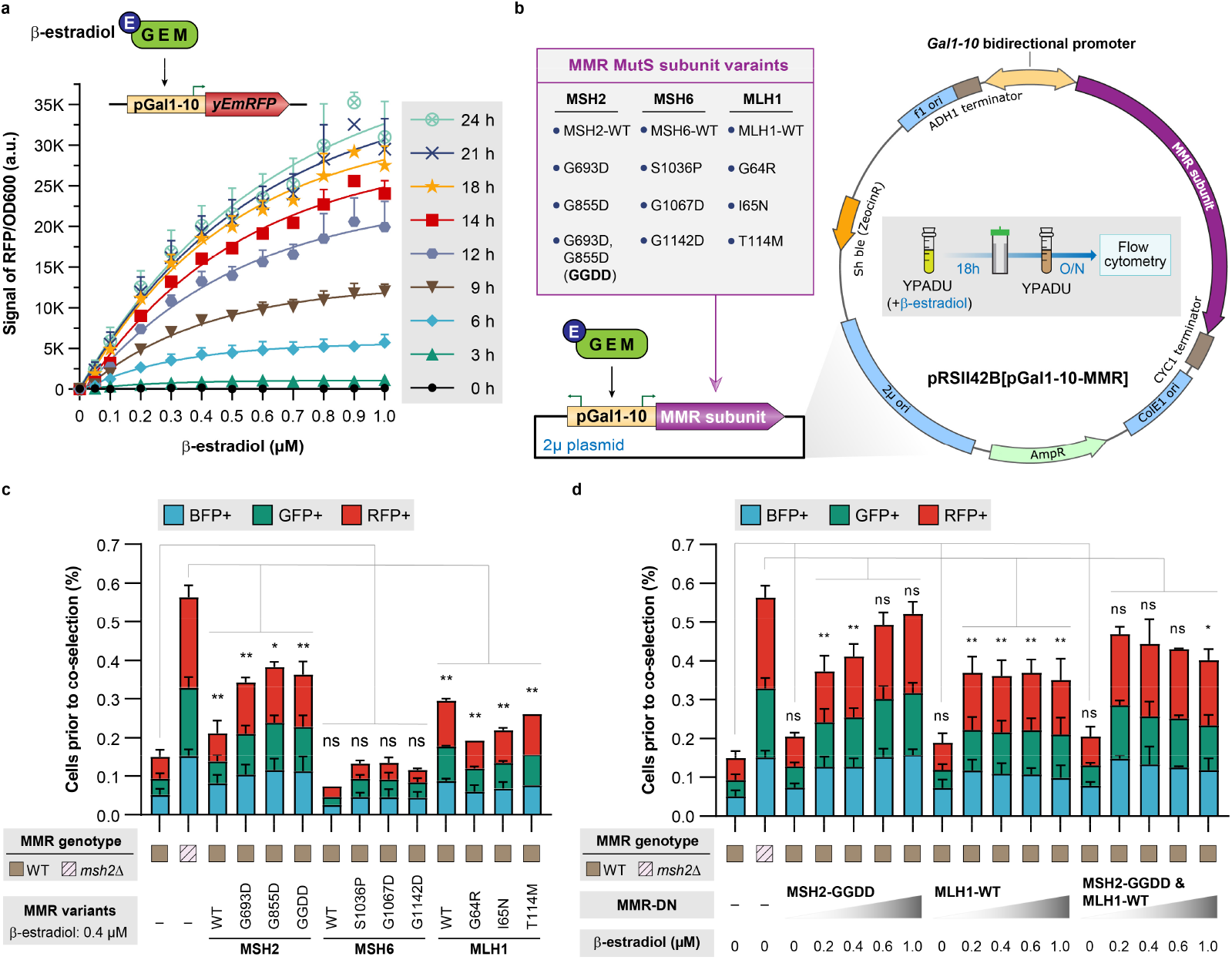
Engineering tunable expression of dominant negative mutants to modulate MMR fidelity during eMAGE editing. **a**, RFP induction over 24 hours with titration of β-estradiol concentration reveals a robust dose-response and time-dependent expression curve. **b**, Selected MMR subunit variants of three wild-type proteins and nine dominant negative mutants overexpressed using our improved β-estradiol inducible system from high-copy 2μ plasmids (see plasmids used in this study in **Supplementary Table 2**). This vector allows inducible expression of up to two MMR subunits simultaneously using the bidirectional *Gal1-10* promoter upon supplying β-estradiol in growth media of strains with GEM expression. Expression of subunit variants from episomal vector was induced with β-estradiol for 18 hours prior to ssODN electroporation. After electroporation cells were recovered overnight in medium without β-estradiol and analyzed by flow cytometry without co-selection. Notably, this vector is also compatible with conventional galactose induction providing flexibility for yeast strains without a pre-integrated GEM cassette. However, galactose induction is not preferable due to the expression level of MMR-DN is more difficult to be controlled and it requires carbon source changes that could perturb cell growth and increase protocol time. **c**, Influence of 12 MMR subunit variants on ssODN-mediated frameshift correction frequency in three fluorescent reporter genes shown in **Fig. 2a**, as compared to MMR-proficient (WT) and MMR-deficient (*msh2*Δ) cells. Expression of subunit variants from episomal vector was induced with 0.4 μM β-estradiol for 18 hours (∼50% of maximum overexpression) prior to ssODN electroporation. **d**, Modulation of MMR fidelity by titrating MSH2-GGDD and MLH1-WT expression with different β-estradiol concentrations. p values of multiple-group comparisons from ordinary one-way ANOVA Dunnett’s test and p values of two-group comparisons from unpaired t-test. ns, not significant, *p < 0.05, **p < 0.01, ***p < 0.001.

## Supplementary Information

**Supplementary Table 1: Yeast Strains used in this study**.

**Supplementary Table 2: Plasmids used in this study**.

**Supplementary Table 3: ssODNs and selective PCR primers used in this study**.

**Supplementary File 1: DNA sequence files of important genomic loci and plasmids**.

## References

1. Chari, R. & Church, G.M. Beyond editing to writing large genomes. Nat Rev Genet 18, 749–760 (2017).

2. Haimovich, A.D., Muir, P. & Isaacs, F.J. Genomes by design. Nat Rev Genet 16, 501–516 (2015).

3. McCarty, N.S., Graham, A.E., Studena, L. & Ledesma-Amaro, R. Multiplexed CRISPR technologies for gene editing and transcriptional regulation. Nature communications 11, 1281 (2020).

4. Komor, A.C., Kim, Y.B., Packer, M.S., Zuris, J.A. & Liu, D.R. Programmable editing of a target base in genomic DNA without double-stranded DNA cleavage. Nature 533, 420–424 (2016).

5. Gaudelli, N.M. et al. Programmable base editing of A*T to G*C in genomic DNA without DNA cleavage. Nature 551, 464–471 (2017).

6. Halperin, S.O. et al. CRISPR-guided DNA polymerases enable diversification of all nucleotides in a tunable window. Nature 560, 248–252 (2018).

7. Strecker, J. et al. RNA-guided DNA insertion with CRISPR-associated transposases. Science 365, 48–53 (2019).

8. Klompe, S.E., Vo, P.L.H., Halpin-Healy, T.S. & Sternberg, S.H. Transposon-encoded CRISPR-Cas systems direct RNA-guided DNA integration. Nature 571, 219–225 (2019).

9. Anzalone, A.V. et al. Search-and-replace genome editing without double-strand breaks or donor DNA. Nature 576, 149–157 (2019).

10. Thompson, D.B. et al. The Future of Multiplexed Eukaryotic Genome Engineering. ACS Chem Biol 13, 313–325 (2018).

11. Barbieri, E.M., Muir, P., Akhuetie-Oni, B.O., Yellman, C.M. & Isaacs, F.J. Precise Editing at DNA Replication Forks Enables Multiplex Genome Engineering in Eukaryotes. Cell 171, 1453–1467 e1413 (2017).

12. Carr, P.A. et al. Enhanced multiplex genome engineering through co-operative oligonucleotide co-selection. Nucleic Acids Res 40, e132 (2012).

13. DiCarlo, J.E. et al. Yeast oligo-mediated genome engineering (YOGE). ACS Synth Biol 2, 741–749 (2013).

14. Costantino, N. & Court, D.L. Enhanced levels of lambda Red-mediated recombinants in mismatch repair mutants. Proceedings of the National Academy of Sciences of the United States of America 100, 15748–15753 (2003).

15. Dekker, M., Brouwers, C. & te Riele, H. Targeted gene modification in mismatch-repair-deficient embryonic stem cells by single-stranded DNA oligonucleotides. Nucleic Acids Res 31, e27 (2003).

16. Nyerges, A. et al. A highly precise and portable genome engineering method allows comparison of mutational effects across bacterial species. Proceedings of the National Academy of Sciences of the United States of America 113, 2502–2507 (2016).

17. Aranda-Diaz, A., Mace, K., Zuleta, I., Harrigan, P. & El-Samad, H. Robust Synthetic Circuits for Two-Dimensional Control of Gene Expression in Yeast. ACS Synth Biol 6, 545–554 (2017).

18. Wu, X.L. et al. Genome-wide landscape of position effects on heterogeneous gene expression in Saccharomyces cerevisiae. Biotechnology for biofuels 10, 189 (2017).

19. Alani, E., Sokolsky, T., Studamire, B., Miret, J.J. & Lahue, R.S. Genetic and biochemical analysis of Msh2p-Msh6p: role of ATP hydrolysis and Msh2p-Msh6p subunit interactions in mismatch base pair recognition. Mol Cell Biol 17, 2436–2447 (1997).

20. Shcherbakova, P.V. & Kunkel, T.A. Mutator phenotypes conferred by MLH1 overexpression and by heterozygosity for mlh1 mutations. Mol Cell Biol 19, 3177–3183 (1999).

21. Das Gupta, R. & Kolodner, R.D. Novel dominant mutations in Saccharomyces cerevisiae MSH6. Nat Genet 24, 53–56 (2000).

22. Lang, G.I. in Genome Instability: Methods and Protocols. (eds. M. Muzi-Falconi & G.W. Brown) 21–31 (Springer New York, New York, NY; 2018).

23. Cao, M., Tran, V.G. & Zhao, H. Unlocking nature’s biosynthetic potential by directed genome evolution. Curr Opin Biotechnol 66, 95–104 (2020).

24. Kaishima, M., Ishii, J., Matsuno, T., Fukuda, N. & Kondo, A. Expression of varied GFPs in Saccharomyces cerevisiae: codon optimization yields stronger than expected expression and fluorescence intensity. Sci Rep 6, 35932 (2016).

25. Keppler-Ross, S., Noffz, C. & Dean, N. A new purple fluorescent color marker for genetic studies in Saccharomyces cerevisiae and Candida albicans. Genetics 179, 705–710 (2008).

26. Subach, O.M., Cranfill, P.J., Davidson, M.W. & Verkhusha, V.V. An enhanced monomeric blue fluorescent protein with the high chemical stability of the chromophore. PLoS One 6, e28674 (2011).

27. Liang, Z., Sunder, S., Nallasivam, S. & Wilson, T.E. Overhang polarity of chromosomal doublestrand breaks impacts kinetics and fidelity of yeast non-homologous end joining. Nucleic Acids Res 44, 2769–2781 (2016).

28. Chee, M.K. & Haase, S.B. New and Redesigned pRS Plasmid Shuttle Vectors for Genetic Manipulation of Saccharomycescerevisiae. G3 2, 515–526 (2012).

29. Brachmann, C.B. et al. Designer deletion strains derived from Saccharomyces cerevisiae S288C: a useful set of strains and plasmids for PCR-mediated gene disruption and other applications. Yeast 14, 115–132 (1998).

30. Gibson, D.G. Enzymatic assembly of overlapping DNA fragments. Methods Enzymol 498, 349–361 (2011).

31. Lang, G.I. & Murray, A.W. Estimating the per-base-pair mutation rate in the yeast Saccharomyces cerevisiae. Genetics 178, 67–82 (2008).

